# Dysbiosis-associated gut bacterium *Ruminococcus gnavus* varies at the strain level in ability to utilize key mucin component sialic acid

**DOI:** 10.1101/2024.08.02.604340

**Authors:** Olga M. Sokolovskaya, Jasmina Uzunovic, Yutian Peng, Mikiko Okumura, Lingjue Mike Wang, Yuhui Zhou, Zijuan Lai, Elizabeth Skippington, Man-Wah Tan

## Abstract

*Ruminococcus gnavus* is a prevalent human gut commensal bacterium with known roles in intestinal mucus degradation, including by catabolism of the terminal mucin sugar sialic acid. While *R. gnavus* is not considered a pathogen, overabundance of this species is correlated with Inflammatory Bowel Disease (IBD), and its sialic acid metabolism may play a role in the dysbiotic state. Interestingly, liberation of mucin-bound sialic acid by *R. gnavus* yields the distinct product of 2,7-anhydro-*N*-acetylneuraminic acid (2,7-anhydro-Neu5Ac), in contrast to other known mucin-degrading bacteria, which generate Neu5Ac. This prompted us to look for 2,7-anhydro-Neu5Ac metabolism proteins in the genomes of 77 *R. gnavus* clinical isolates. We found that 2,7-anhydro-Neu5Ac metabolism is sporadically distributed in this species with respect to phylogeny and strain origin. We measured sialic acid-dependent growth of 12 sequenced isolates, finding that the presence of 2,7-anhydro-Neu5Ac catabolism proteins was predictive of growth on this substrate. Our analysis also uncovered “partial” 2,7-anhydro-Neu5Ac catabolism pathways in two *R. gnavus* strains, which we determined constitute the canonical Neu5Ac catabolism pathway, previously unreported in this species. These results reveal a notable diversity of sialic acid catabolism across the *R. gnavus* species, an essential consideration for further investigations into the importance of this metabolism in mucin degradation and in roles of *R. gnavus* in IBD and other gut dysbioses.

## Introduction

*Ruminococcus (Mediterraneibacter) gnavus*^1^ is a commensal human gut bacterium that has come into the spotlight in the microbiome field due to its association with disease. While it is a prevalent gut bacterium, found in over 90% of individuals, multiple studies link *R. gnavus* overabundance with Inflammatory Bowel Disease (IBD)^2–12^. Despite reported associations, it remains unknown whether *R. gnavus* becomes abundant as a result of inflammation (for example, by higher tolerance of increased oxygen concentrations associated with the inflamed state, compared to other anaerobic gut bacteria^2^), or whether it exacerbates or plays causative roles in disease (for example, by surface presentation of ‘superantigens’^13^, capsular polysaccharide^14^, or production of pro-inflammatory factors^15^). Importantly, the *R. gnavus* species contains significant strain-level variation, according to a pangenome analysis of 17 *R. gnavus* strains that reports 1,708-2,473 variable genes per genome, compared to a core genome of 1,178 genes^2^. The strain-level variation in this organism, combined with the fact that individuals can harbor multiple strains, complicates questions about the potential roles of *R. gnavus* in health and disease.

A genomic analysis of *R. gnavus* strains published by Hall *et al.* reports that a sialic acid transporter is among 199 proteins found exclusively within a clade of *R. gnavus* that is comprised of IBD-associated strains, suggesting the relevance of sialic acid metabolism in the diseased state^2^. In the colon, the region of the gut that bares the highest microbial load, sialic acids, particularly *N*-acetylneuraminic acid (Neu5Ac), are abundant terminal sugars on glycosylated mucins, the main constituents of the mucus lining of the gut^16^. Because it caps glycan chains, Neu5Ac plays a role in mucus integrity, protecting mucins from degradation by bacterial glycoside hydrolases and mucinases^17^. This role of Neu5Ac is notable, as mucus degradation is a hallmark of IBD^18–22^. Simultaneously, specialized mucin degrading microorganisms express sialidases that liberate glycan-linked Neu5Ac, and free Neu5Ac is an important growth substrate for certain commensal gut bacteria and pathogens^23–28^. Because Neu5Ac released by extracellular sialidases is cross-fed between different bacterial species, Neu5Ac plays important ecological roles in the gut environment^25–27,29,30^. In fact, Neu5Ac cross-feeding enables the expansion of opportunistic pathogens in mouse models of colitis^26,27^.

Interestingly, *R. gnavus* cleaves sialic acid by a mechanism that is distinct from known mucin-degrading bacteria, namely by releasing 2,7-anhydro-Neu5Ac, rather than free Neu5Ac, as the cleavage product of glycan-linked Neu5Ac^31^. This reaction is catalyzed by the NanH enzyme, an intramolecular *trans* (IT)-sialidase. Notably, IT domains were reported to be enriched in the metagenomes of individuals with IBD compared to healthy controls^31^. *R. gnavus* is able to use 2,7-anhydro-Neu5Ac as a growth substrate by virtue of an ABC family transporter that selectively imports 2,7-anhydro-Neu5Ac (but not Neu5Ac) into the cell^24^, and an oxidoreductase NanOx which converts internalized 2,7-anhdyro-Neu5Ac into Neu5Ac^24^, which is further metabolized by the canonical Neu5Ac catabolism enzymes NanA, NanE, and NanK (Figure 1A). The ABC transporter and *nan* genes are found within a single cluster in the *R. gnavus* genome^24^. Because most sialic acid-metabolizing bacteria cannot consume 2,7-anhydro-Neu5Ac, this metabolism by *R. gnavus* is considered a “selfish” behavior.

**Figure 1.**
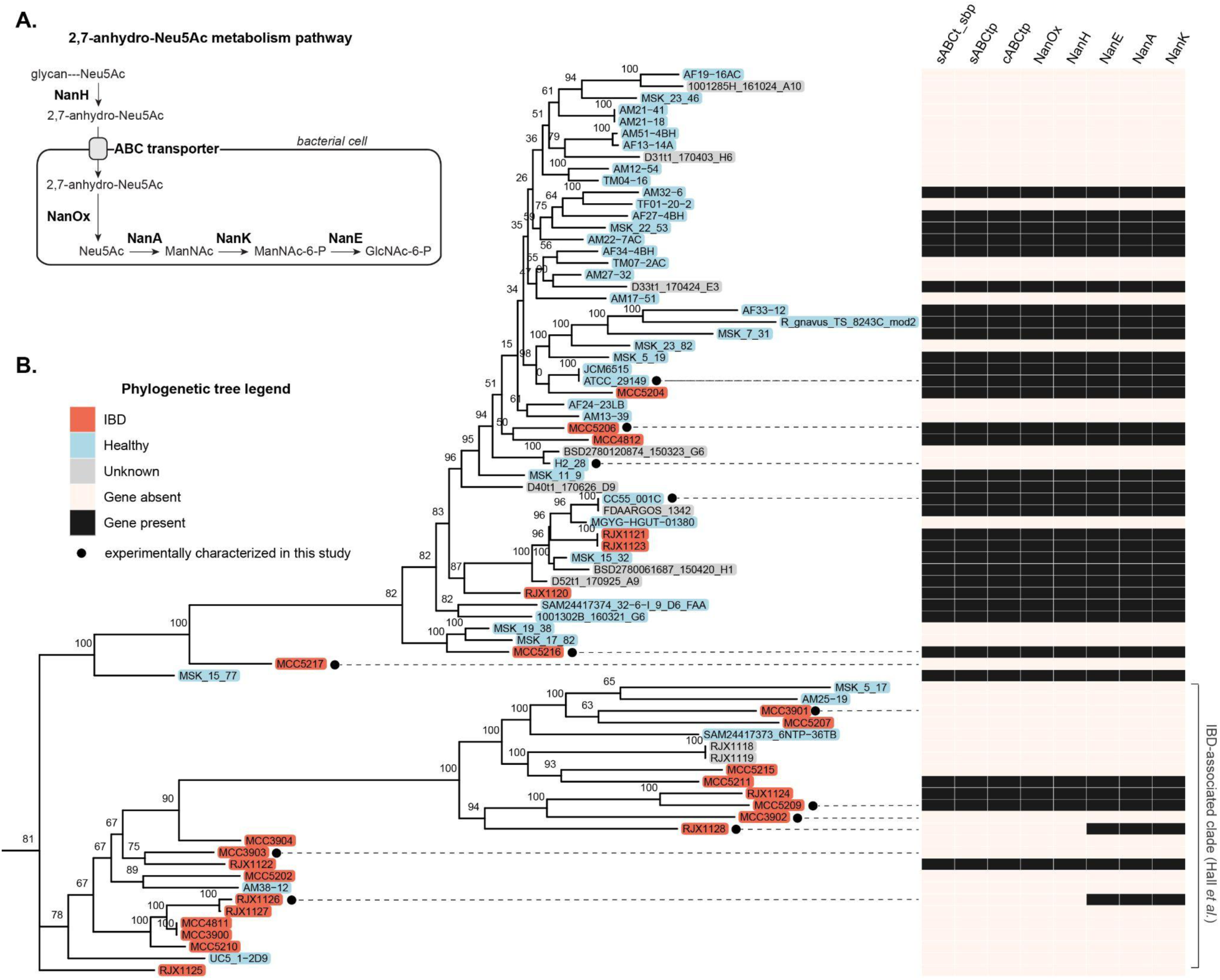
(A) Proteins involved in 2,7-anhydro-Neu5Ac catabolism in *R. gnavus*. Abbreviated metabolites: ManNAc, *N*-acetylmannosamine; ManNAc-6-P, *N*-acetylmannosamine-6-phosphate; GlcNAc-6-P, *N*-acetylglucosamine-6-phosphate. (B) *R. gnavus* strain-level phylogeny and evidence for Nan protein-gene content for 77 sequenced *R. gnavus* genomes. The phylogenetic tree is based on a maximum-likelihood analysis of a 1,415,350 position core gene alignment, and is unrooted. Branch labels indicate bootstrap support from 500 replicates. Strains names are colored based on their origins. Shown on the right is a genome-wide BLAST comparison of Nan protein-coding sequences. Filled boxes indicate high-scoring segment pairs (HSPs) yielded by BLASTp searches of putative protein sequences for the corresponding query *R. gnavus* genome against Nan protein sequences from reference strain *R. gnavus* ATCC 29149 with default parameter settings. Only HSPs satisfying > 80% sequence identity and > 60% sequence coverage are shown. Protein annotations: sABCt_sbp, sugar ABC transporter substrate binding protein; cABCtp, carbohydrate ABC transporter permease; sABCtp, sugar ABC transporter permease.

The aforementioned studies implicating *R. gnavus* sialic acid metabolism in disease, together with the unique mechanism of sialic acid cleavage by this organism, and limited data revealing that sialic acid metabolism and growth on mucin vary between *R. gnavus* strains^32,33^, prompted us to look more specifically into the distribution of sialic acid catabolism across clinical isolates of *R. gnavus*. By analyzing 77 *R. gnavus* genomes, including 17 newly sequenced in this study, from healthy people as well as individuals with IBD, we were able to profile the distribution of sialic acid catabolism within the *R. gnavus* species, and discovered previously unknown variation in sialic acid utilization. Our findings underscore the essentiality of considering specific strains in future investigations into the roles of *R. gnavus* sialic acid metabolism in the gut.

## Results

### Sialic acid metabolism is sporadically distributed among R. gnavus strains

The prevalence of 2,7-anhydro-Neu5Ac catabolism within the *R. gnavus* clade was investigated by comparing protein sequences from 77 *R. gnavus* genomes to Nan protein sequences from reference strain ATCC 29149. The 77 genomes comprise 60 that were publicly available at the time the analysis commenced, and a further 17 that were sequenced from an in-house collection, to which we refer as “MCC” strains. We included a significant proportion of strains from healthy individuals (41 of 77) to address a limitation of previous strain-level genomic analyses of *R. gnavus*, which have predominantly focused on isolates from patients with IBD. To investigate the Hall *et al*. finding that a sialic acid transporter may be constrained to a particular clade of *R. gnavus*^2^, we constructed a core genome phylogenetic tree of *R. gnavus* strains and indicated the putative presence or absence of the 2,7-anhydro-Neu5Ac catabolism proteins across the strain tree (Figure 1B).

Like Hall *et al*., we observed two clades of *R. gnavus*, with one clade showing a higher proportion of, but not exclusively composed of, IBD-associated strains (Figure 1B; IBD-associated clade is marked on the far right). We found that roughly half (35/77) of the genomes encoded protein sequences with homology to Nan proteins, including strains isolated from healthy individuals (17/41 with Nan proteins) and from patients with IBD (11/26 with Nan proteins). All but two of these sequences yielded BLAST high scoring segment pairs (HSPs) for all of the proteins in the 2,7-anhydro-Neu5Ac catabolism pathway, namely the three-component ABC transporter (a substrate binding protein and two transporter permeases), NanH, NanOx, NanE, NanA, and NanK (Figure 1B). Among four divergent strains we found ≥ 98.9% sequence identity in the Nan proteins at the amino acid level, and apparent synteny of the *nan* gene cluster (Figure S1). Strains RJX1126 and RJX1128 had “partial” pathways, yielding HSPs for NanE, NanA, and NanK only (Figure 1B), which we examine in further detail in a following section. Overall, our analysis revealed that the 2,7-anhydro-Neu5Ac catabolism pathway is sporadically distributed throughout phylogenetically diverse strains. Thus, the sialic acid transporter that was identified by Hall *et al*. as belonging exclusively to an IBD-associated *R. gnavus* clade is unlikely to be the ABC family sialic acid transporter previously reported in this species.

### Sialic acid-dependent growth is consistent with the presence or absence of Nan proteins

To determine how the presence of Nan proteins corresponds to sialic acid utilization by live bacterial cells, we selected a dozen strains that span the phylogenetic diversity of this species (marked in Figure 1B and listed in Table S1) and cultured them in basal YCFA medium supplemented with 2,7-anhydro-Neu5Ac, Neu5Ac, and 3’-sialyllactose, a human milk oligosaccharide consisting of sialic acid linked to lactose. Like *R. gnavus* ATCC 29149, all strains whose genomes encode all five Nan proteins and an ABC family transporter grew on 2,7-anhydro-Neu5Ac and 3’-sialyllactose, but not Neu5Ac (Figure 2A). The absence of growth of all but one of these strains on lactose (Figure S2) provides evidence that the strains use the sialic acid component of 3’-sialyllactose as a growth substrate. No major differences in growth rate were observed between strains grown with 2,7-anhydro-Neu5Ac or 3’-sialyllactose (Figure S3). Strains lacking Nan proteins did not grow on any source of sialic acid (Figure 2A).

**Figure 2.**
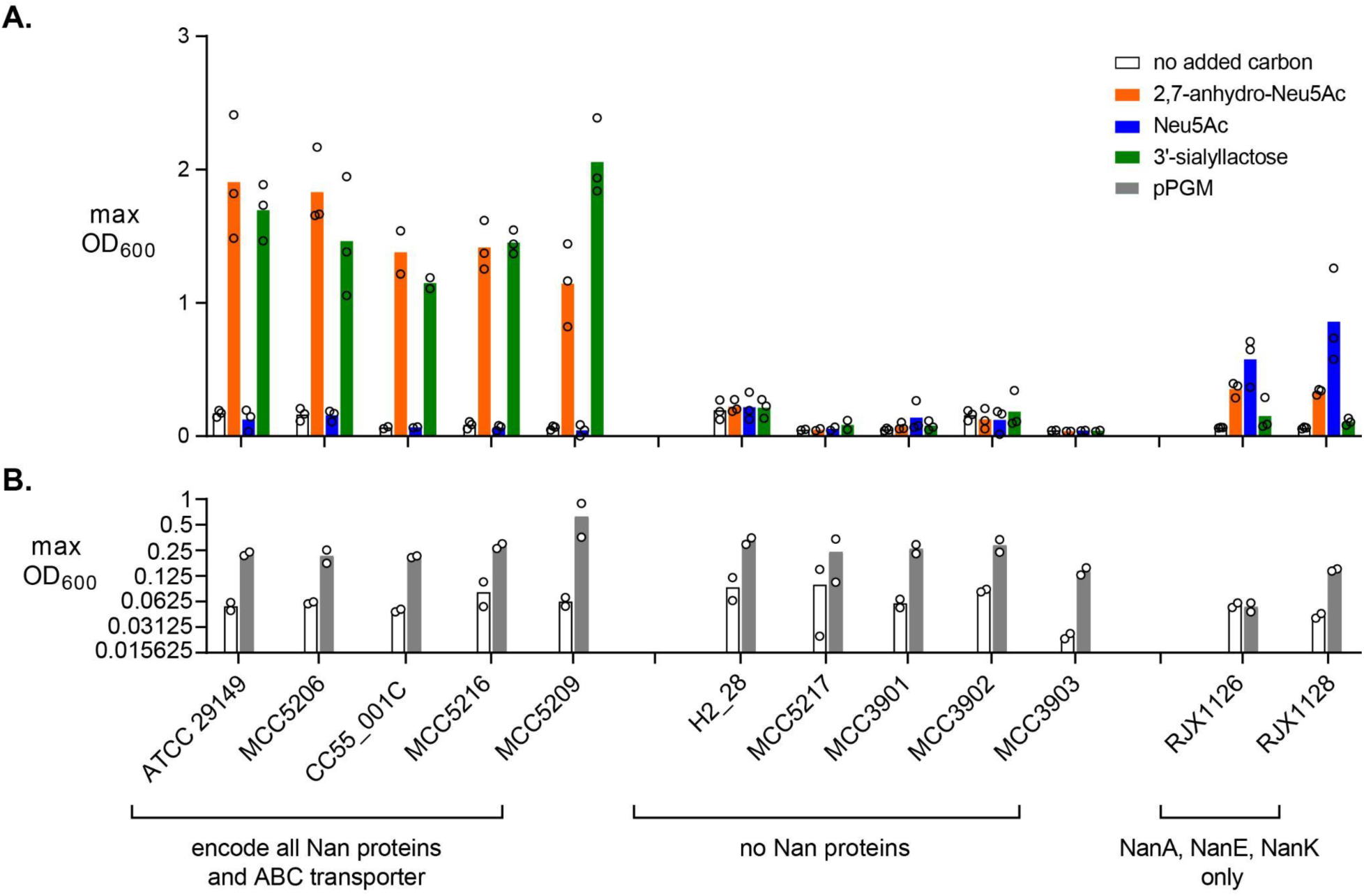
Growth of *R. gnavus* strains on various carbon sources. Maximum optical densities reached by strains in culture during 24 hours of growth on (A) simple sugars or (B) purified porcine gastric mucin (pPGM). Bars indicate the average of at least two independent experiments, each of which is represented by a data point.

Unlike the aforementioned strains, RJX1126 and RJX1128 grew to the highest density with Neu5Ac, and were unable to grow with 3’-sialyllactose (Figure 2A). This phenotype is consistent with the presence of NanA, NanK, and NanE enzymes, and the absence of the sialidase NanH. Both strains also achieved an intermediate density when grown with 2,7-anhydro-Neu5Ac. The apparent stability of 2,7-anhydro-Neu5Ac in bYCFA over the course of 24 hours (Figure S4) suggests that growth was not due to spontaneous conversion of 2,7-anhydro-Neu5Ac to Neu5Ac, although an indirect effect of bacterial growth on 2,7-anhydro-Neu5Ac stability cannot be ruled out.

Finally, we screened all 12 *R. gnavus* clinical isolates for the ability to grow with purified porcine gastric mucin (pPGM). As previously reported, strain ATCC 29149 reached a moderate density with pPGM as the sole added carbon source^32^. Notably, all of the strains except for RJX1126 grew with pPGM, doubling two to three times over background (Figure 2B), with some variation in growth dynamics (Figure S5). Given that seven of these 12 strains were unable to grow with 3’-sialyllactose (i.e. cannot utilize glycan-linked sialic acid as a growth substrate), it is clear that sialic acid catabolism is not a sole determinant of *R. gnavus* growth on pPGM.

### The Nan protein cluster in *R. gnavus* strains RJX1126 and RJX1128 functions in Neu5Ac catabolism

The NanA, NanE, and NanK proteins found in *R. gnavus* strains RJX1126 and RJX1128 appear to be distinct homologs from those found in other *R. gnavus* strains; these proteins share 91.5%, 86.96%, and 80.82% amino acid identity, respectively, with the ATCC 29149 homologs, whereas the NanA, NanE, and NanK sequences in all other strains are more highly conserved (Figure S1). Furthermore, the *nanAEK* genes in *R. gnavus* strains RJX1126 and RJX1128 are found within a distinct gene cluster (Figure 3A and Table 1) that lacks the 2,7-anhydro-Neu5Ac metabolism-specific genes *nanH* and *nanOx*, and lacks the ABC family 2,7-anhydro-Neu5Ac importer. Instead, the *nan* genes in these two strains are clustered with a sodium:solute symporter (SSS) gene. SSS family transporters are one of four classes of transporters involved in the import of Neu5Ac^34^, and the SSS protein in *R. gnavus* shares 68.6% identity with a verified Neu5Ac transporter protein in *Clostridioides (Clostridium) difficle*^26^. Interestingly, this locus also encodes proteins putatively involved in transposition and reverse transcription, suggesting it may have been horizontally acquired.

**Figure 3.**
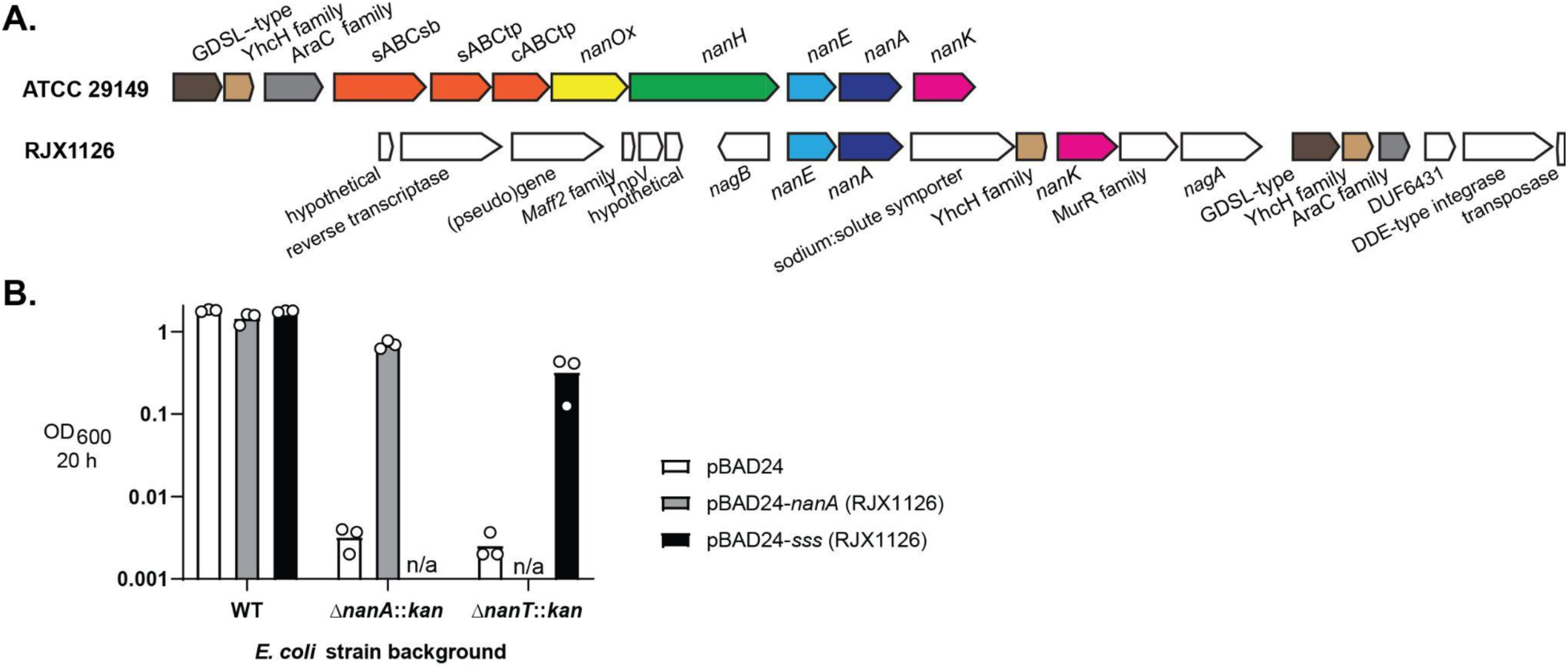
(A) Gene neighborhood of the *nan* genes in *R. gnavus* strain RJX1126, contrasted to the previously described *nan* cluster in strain ATCC 29149. (B) Complementation of *E. coli* BW25113 *nanA* and *nanT* (sialic acid transporter) mutants with *R. gnavus* RJX1126 *nanA* and sodium:solute symporter (*sss*) genes, tested by growth in M9 medium with Neu5Ac as the sole carbon source. Data points show the density achieved by each strain after 20 h of growth, in three independent experiments; columns indicate mean OD_600_ values. “n/a”: this condition was not tested.

**Table 1.** Putative sialic acid catabolism gene cluster in *R. gnavus* strain RJX1126 (genome accession number NZ_NIHU01000030.1)

| Locus Tag | Genomic location | Current NCBI annotation | Comments |
| --- | --- | --- | --- |
| CDL24_RS14640 | 1-132 | transposase domain-containing protein |  |
| CDL24_RS14645 | 178-1497 | DDE-type integrase/transposase/recombinase |  |
| CDL24_RS14650 | 1571-2041 | DUF6431 domain-containing |  |
| CDL24_RS14655 | 2242-2697 | AraC family ligand binding domain-containing protein | transcriptional regular |
| CDL24_RS14660 | 2768-3223 | YhcH/YjgK/YiaL family protein | possible sugar anomerase |
| CDL24_RS14665 | 3244-3954 | GDSL-type esterase/lipase family protein | possible acyl hydrolase |
| CDL24_RS14670 | 4388-5542 | N-acetylglucosamine-6-phosphate deacetylase | *nagA* |
| CDL24_RS14675 | 5587-6426 | MurR/PriR family transcriptional regulator |  |
| CDL24_RS14680 | 6445-7332 | ROK family protein | putative *nanK* |
| CDL24_RS14685 | 7467-7919 | YhcH/YjgK/YiaL family protein | possible sugar anomerase |
| CDL24_RS14690 | 7935-9440 | sodium:solute symporter | putative Neu5Ac transporter |
| CDL24_RS14695 | 9546-10463 | dihydrodipicolinate synthase family protein | putative *nanA* |
| CDL24_RS14700 | 10518-11207 | N-acetylmannosamine-6-phosphate 2-epimerase | *nanE* |
| CDL24_RS14705 | 11488-12216 | glucosamine-6-phosphate deaminase | *nagB* |
| CDL24_RS14715 | 12710-12991 | hypothetical protein |  |
| CDL24_RS14720 | 13006-13353 | TnpV protein | transposon encoded protein |
| CDL24_RS14725 | 13427-13588 | Maff2 family protein | short transmembrane protein |
| CDL24_RS14730 | 13865-15187 | uncharacterized (pseudo)gene |  |
| CDL24_RS14735 | 15297-16787 | reverse transcriptase domain-containing protein |  |
| CDL24_RS15960 | 16903-17103 | hypothetical protein |  |

The genetic content of the *nan* cluster in *R. gnavus* strains RJX1126 and RJX1128 is consistent with the ability of these two strains to utilize Neu5Ac as a growth substrate (Figure 2A). To probe the hypothesis that the distinct gene cluster we discovered in these strains functions in Neu5Ac catabolism, we tested whether the RJX1126 *nanA* and SSS-encoding genes would complement mutations in the Neu5Ac catabolism pathway of the laboratory strain *Escherichia coli* BW25113. The RJX1126 *nanA* gene restored Neu5Ac-dependent growth of *E. coli* containing a *nanA* disruption, and the SSS gene restored the growth of an *E. coli nanT* (Neu5Ac transporter of the major facilitator superfamily) mutant, demonstrating that these *R. gnavus* RJX1126 genes indeed function in Neu5Ac import and catabolism. The presence of the SSS in *R. gnavus* strains RJX1126 and RJX1128 likely confers their unique (within the species *R. gnavus*) ability to grow with Neu5Ac, as the SSS protein is absent from all other *R. gnavus* genomes examined in this study.

## Discussion

The disease-associated, gut commensal species *R. gnavus* catabolizes sialic acid by a mechanism that differs from most sialic acid-metabolizing organisms^31^. Because of the known roles of sialic acid catabolism in mucus homeostasis and microbial ecology^17,26,27,29^, and due to multiple reports that *R. gnavus* is positively associated with gut inflammatory disease^2–12^, the biological impacts of the 2,7-anhydro-Neu5Ac metabolism of this organism were of interest to us. Indeed, sialic acid and mucus metabolism are known to vary between strains of *R. gnavus*, but, prior to this research, few *R. gnavus* strains had been investigated in this regard^32,33^. Here, we surveyed sialic acid catabolism across 77 strains, finding that 2,7-anhydro-Neu5Ac catabolism was sporadically distributed among strains isolated from both healthy individuals and patients with IBD, and discovering further diversity than was previously appreciated. Based on our results, we can classify *R. gnavus* strains into three categories of sialic acid metabolism: strains that cleave glycan-linked sialic acid to generate 2,7-anhydro-Neu5Ac and use this substrate for growth; strains that require free Neu5Ac for sialic acid-dependent growth; and strains that do not utilize sialic acid as a growth substrate (Figure 4).

**Figure 4.**
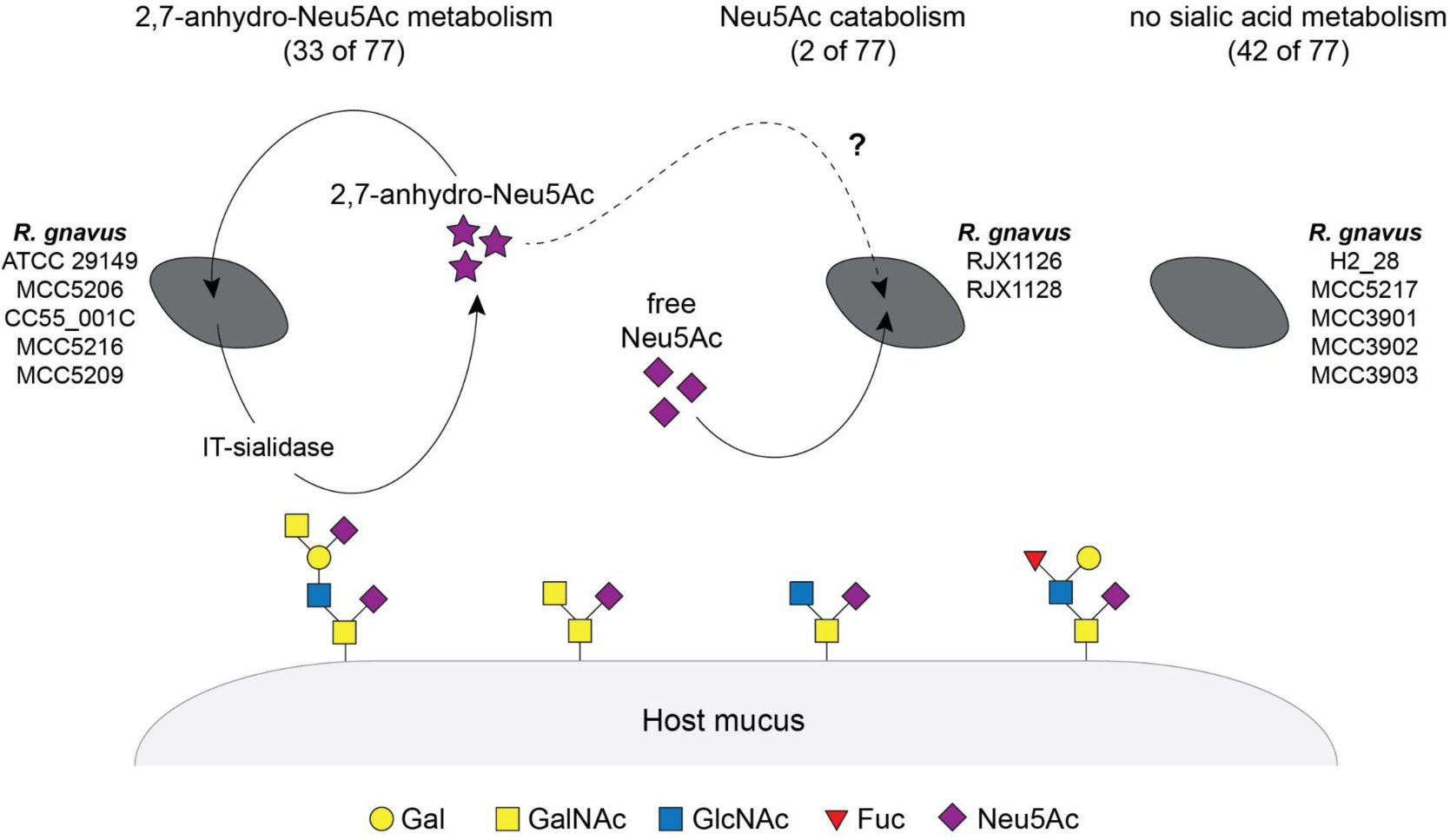
Variable interactions of *R. gnavus* strains with mucus-derived sialic acids. Left: strains that cleave glycan-linked Neu5Ac to generate 2,7-anhydro-Neu5Ac, and use 2,7-anhydro-Neu5Ac as a growth substrate. Middle: strains that grow with free Neu5Ac (and to a lesser extent with free 2,7-anhydro-Neu5Ac, although this phenotype requires further investigation). Right: strains that do not catabolize sialic acid. The numbers above each cartoon indicate the fraction of analyzed *R. gnavus* genomes that fall into each category of sialic acid metabolism; strains that were experimentally characterized in this work are listed in their corresponding category. Glycans are depicted according to Symbol Nomenclature for Glycan Systems, and represent abundant sialylated oligosaccharides in the human colonic mucin MUC2^38,39^ (Gal, galactose; GalNAc, *N*-acetylgalactosamine; GlcNAc, *N*-acetylglucosamine; Fuc, fucose).

As mentioned in describing the motivation for this study, Hall *et al.* identified a sialic acid transporter that was absent from isolates from healthy individuals and restricted to an IBD-specific clade^2^. Based on our examination of the distribution of sialic acid catabolism across *R. gnavus* strains, we conclude that the sialic acid transporter that emerged from their study was the Neu5Ac SSS family transporter that we identified in strains RJX1126 and RJX1128. The potential relevance of Neu5Ac catabolism for *R. gnavus* growth in inflammatory conditions is of interest given that high free Neu5Ac levels are associated with dysbiosis^26,27^. Curiously, strains RJX1126 and RJX1128 also grew with 2,7-anhydro-Neu5Ac as the sole carbon source, albeit to a significantly lower density compared to strains that have the complete *nan* cluster. As the genomes of strains RJX1126 and RJX1128 lack the gene encoding NanOx, it is unclear whether their growth on 2,7-anhydro-Neu5Ac is due to enzymatic conversion of this substrate to Neu5Ac or a non-specific process. However, there is precedent for substrate promiscuity in sialic acid transporters^34,35^, so future experiments testing the ability of the *R. gnavus* SSS family transporter to import 2,7-anhydro-Neu5Ac in addition to Neu5Ac will help elucidate the mechanism of growth on this substrate.

Despite variations in sialic acid catabolism among the *R. gnavus* strains we tested, all but one of the strains grew with pPGM as the sole carbon source. Importantly, this finding may or may not translate to growth on human intestinal mucin, which is expected to differ significantly in glycan composition compared to PGM. Previous work on established mucin degraders demonstrates that growth on oligosaccharides derived from different mucin sources can vary significantly, for example depending on the degree of glycan sulfation^36^. However, taken at face value, the finding that almost all of the *R. gnavus* strains tested grew on mucin suggests that different strains have evolved different mechanisms to utilize this important nutrient source. A previous study that compared the repertoire of glycoside hydrolases in *R. gnavus* strain ATCC 29149 with those present in strain E1, which is unable to grow on pPGM, hypothesized that specific α-L-fucosidases may play a role in the ability of *R. gnavus* ATCC 29149 to grow on PGM^32^. Like sialidase expression, fucosidase expression, as well as endo-galactosidase expression, was upregulated in *R. gnavus* strains in the presence of mucin^32,33^. Similar comparative genomics approaches across a larger number of *R. gnavus* strains may be useful in generating testable hypotheses regarding mechanisms of growth on mucin.

The biological impact of differential sialic acid catabolism in *R. gnavus* is an exciting topic of future exploration. The finding that *R. gnavus* strains have evolved different ways to utilize sialic acid suggests the importance of this nutrient source in the gut environment. Based on the distinct sialic acid catabolism pathways presented in Figure 4, we hypothesize that different *R. gnavus* strains would differentially respond to changes in mucus (e.g. quantity of mucus, degree of mucin sialylation) and would be differentially impacted by levels of general sialidase activity, and consequently different levels of free sialic acid, which are known to be altered in healthy versus diseased states of the gut^26,27^. Additionally, we would predict distinct ecological interactions of *R. gnavus* with other gut bacteria based on sialic acid catabolism. For example, *R. gnavus* strains that generate 2,7-anhydro-Neu5Ac may be involved in cross-feeding interactions with other 2,7-anhydro-Neu5Ac-metabolizing bacteria and may effectively sequester sialic acid from bacteria that typically take advantage of Neu5Ac. In contrast, *R. gnavus* strains that can utilize Neu5Ac as a growth substrate might benefit from the presence of gut bacterial species that are Neu5Ac producers. These types of interactions may contribute to changes in microbial community composition that occur in gut dysbiosis, which is a critical component of IBD^37^. Thus, examining specific and relevant *R. gnavus* strains will be key to untangling potential roles of the sialic acid metabolism of this species in gut biology in health and disease.

## Materials and Methods

### Materials

2,7-anhydro-Neu5Ac was prepared from 2ʹ-(4-methylumbelliferyl)-*α*-D-*N*-acetylneuraminic acid (4-MU-Neu5Ac) purchased from Carbosynth (see “Enzymatic synthesis of 2,7-anhydro-Neu5Ac”). *N*-acetylneuraminic acid was purchased from Sigma and Carbosynth. 3’-sialyllactose was purchased from Carbosynth. Porcine gastric mucin (type III) was purchased from Sigma and further purified (see “Mucin preparation”). Sources of *R. gnavus* strains are detailed in Supplementary Table 1^2,40–43^.

### Strain isolation and genome sequencing

*R. gnavus* strains were isolated from stool samples from participants of phase 3 trials for the treatment of Crohn’s disease and Ulcerative Colitis^42,43^. Isolations were performed by Microbiotica Ltd using a proprietary platform that utilizes sequencing- and culturing-guided strain selection^44^. Genomic DNA was extracted using the Qiagen DNeasy UltraClean Microbial Kit. Quality of the genomic DNA was assessed using the Genomic DNA ScreenTape and TapeStation 4200 (Agilent Technologies). Genomic DNA was quantified using the Qubit dsDNA BR assay kit (Thermo Fisher Scientific). For library preparation, the Nextera DNA Flex kit (Illumina) was used with an input of 100 ng of genomic DNA. The resulting libraries were multiplexed and sequenced on NovaSeq (Illumina) to generate 5 million paired end 75 base pair reads for each sample. Genomes were assembled and annotated with Spades v3.13.1^45^ and Prokka v1.12^46^, respectively, using default settings.

### Phylogenetic tree construction and annotation

An *R. gnavus* phylogenetic tree was constructed using 60 publically available genomes downloaded from the NCBI GenBank database and 17 genomes sequenced in this study (see “Strain isolation and genome sequencing”). Roary^47^ was used to construct a core gene alignment (present in 95% of isolates), from which a phylogenetic tree was estimated with RAxML v8^48^ using the generalized time-reversible (GTR) model with gamma correction for among-site rate variation. Support for nodes was assessed using 500 bootstrap replicates. The tree is unrooted. R packages tidyverse, ggtree, ape, treeio were used for visualization. Strain origins were annotated by referencing source studies. Genome protein sequences were compared by BLASTp v2.2.28^49^ to Nan protein sequences from *R. gnavus* ATCC 29149. A protein was considered present if BLAST searches yielded HSPs satisfying > 80% identity and > 60% sequence coverage.

### Bacterial culturing

*R. gnavus* was cultured inside of a Coy anaerobic chamber, with a headspace of 85% N_2_/5% H_2_/10% CO_2_, in basal YCFA (bYCFA; containing, per liter: 10 g casitone, 2.5 g yeast extract, 4 g NaHCO_3_, 1 g L-cysteine, 450 mg K_2_HPO_4_, 450 mg KH_2_PO_4_, 900 mg (NH_4_)_2_SO_4_, 900 mg NaCl, 90 mg MgSO_4_, 90 mg CaCl_2_, 1 mg resazurin, 10 mg hemin, 10 µg biotin, 10 µg cyanocobalamin, 30 µg *p*-aminobenzoic acid, 50 µg folic acid, 150 µg pyridoxamine, 4.05 mL acetate, 1.43 mL propionate, 0.22 mL n-valerate, isovalerate and isobutyrate, pH 6.5; and 50 µg thiamine, 50 µg riboflavin, added after autoclaving) supplemented with 0.2% glucose, at 37 °C. For testing growth on various carbon sources, *R. gnavus* strains were pre-cultured overnight in bYCFA with 0.1% glucose and diluted 1:200 into fresh bYCFA containing the specified growth substrate at a concentration of 11.1 mM, or 1 % w/v mucin. Growth rates were measured by culturing in 96-well plates in an Epoch2 plate reader (BioTek), at 37 °C, with continuous orbital shaking.

### Heterologous expression and purification of NanH for 2,7-anhydro-Neu5Ac synthesis

The *R. gnavus* ATCC 29149 *nanH* gene (truncated to remove the N-terminal signaling sequence) was codon optimized for *E. coli* and cloned with an N-terminal hexahistidine tag into the pET11a vector (see Supplementary Materials and Methods) for expression in *E. coli* BL21-Gold(DE3). The expression strain carrying the pET11a-*nanH* plasmid was pre-cultured at 37 °C in LB medium with carbenicillin (50 ug/mL) until OD_600_ 0.8-0.9, after which protein expression was induced at 16 °C for 24 hours by addition of 1 mM isopropyl β-D-1-thiogalactopyranoside. Frozen bacterial cell pellets were resuspended in 50 mM Tris-HCl, 150 mM sodium chloride, pH 7.8, supplemented with 10 mM imidazole, cOmplete Mini EDTA-free Protease Inhibitor Cocktail, and 0.1 mg/mL DNase, and lysed with BugBuster Protein Extraction Reagent (Millipore). The protein was batch bound to Ni-NTA resin, washed with 50 mM Tris-HCl, 150 mM sodium chloride, pH 7.8, 10 mM imidazole, and eluted using an increasing concentration of imidazole in the same buffer. Eluate purity was analyzed by SDS-PAGE using Coomassie stain, and fractions enriched in NanH were concentrated and exchanged into 10 mM HEPES, pH 8.0, with 10% glycerol by repeated centrifugation in Amicon 30 kDa molecular weight cut off (MWCO) spin filters. Purified protein was flash frozen with liquid nitrogen and stored at -80 °C.

### Enzymatic synthesis of 2,7-anhydro-Neu5Ac

2,7-anhydro-Neu5Ac was prepared by treatment of 4-MU-Neu5Ac (100 mM) with *R. gnavus* NanH enzyme (0.01 mg/mL) in 20 mM Na_2_HPO_4_, 20 mM NaH_2_PO_4_, pH 6.5, containing 1 mg/ml bovine serum albumin, at 37 °C. The progression of the reaction was monitored fluorimetrically (excitation 340 nm, emission 420 nm). Upon completion of the reaction, the leaving group 4-methylumbelliferone crystalized out of solution and was removed. The reaction was then quenched by addition of ethanol (50% v/v) and precipitate was removed by filtration. An Amicon 10 MWCO spin filter was used to remove residual protein, and product purity was evaluated by liquid chromatography-mass spectrometry (LC-MS) and 1D/2D ^1^H NMR (see Supplementary Materials and Methods). 2,7-anhydro-Neu5Ac was dried, quantified using quantitative NMR (with caffeine as an external standard), dissolved in water at a stock concentration of 222 mM, and stored at -20° C.

### Mucin preparation

Mucin was dissolved in Dulbrecco’s phosphate buffered saline, pH 7.4. After 1-2 hours stirring, the pH was adjusted to 7.4 and the suspension was stirred overnight. Insoluble debris was removed by centrifugation (10000 x g, 4 degrees, 30 minutes). Mucin was further purified and sterilized by addition of ice-cold ethanol (60% v/v, final). The precipitated mucin was collected by gentle centrifugation (250 x g, 1 minute) and resuspended in water. To remove residual salts, the mucin solution was dialyzed against water using a 10 MWCO Spin-A-Lyzer G2 dialysis cassette (Pierce Biotechnology). Purified mucin was lyophilized and stored at -20 °C.

### Genetic complementation assays

Predicted *nanA* and sodium:solute symporter (*sss*) open reading frames (contig NIHU0100030.1, nucleotides 7000..7917 and 8023..9528, respectively) were amplified from *R. gnavus* strain RJX1126 and cloned into pBAD24 at the NcoI restriction site. Plasmids were electroporated into *E. coli* Keio collection strains^50^ (wildtype, Δ*nanA*::*kan*, and Δ*nanT*::*kan*). Transformed strains were grown in M9 medium with 0.2% glucose and 50 µg/mL carbenicillin, and transferred to M9 medium containing 11.1 mM Neu5Ac and 50 µg/mL carbenicillin for growth assays. Because of leaky expression of the arabinose-inducible promoter, complementation was achieved in the absence of inducer (addition of arabinose was toxic to strains expressing *R. gnavus nanA* or *sss*; data not shown). Growth kinetics were measured by culturing in a 96-well plate in a BioTek Synergy2 plate reader at 37 °C with continuous shaking.

## Data availability statement

Sequencing data are available from NCBI under BioProject ID PRJNAXXXXXXX with BioSample accessions SAMNXXXXXXXX-XX.

## Acknowledgements

We would like to thank Fiona Tamburini for generating early versions of the *R. gnavus* strain-level phylogenetic tree; Genentech colleagues and Kristopher Kennedy for many helpful discussions about this work; and Dr. Ramnik Xavier and his laboratory members for generously sharing strains RJX1126 and RJX1128.

---

**Supplementary Table 1.** Bacterial *s*trains used in this study.

| Strain name | Species | Source |
| --- | --- | --- |
| ATCC 29149 | *Ruminococcus gnavus* | American Type Culture Collection (ATCC)<sup>1</sup> |
| CC55_001C (HM-1056) | *Ruminococcus gnavus* | BEI Resources<sup>2</sup> |
| H2_28 (DSM108212) | *Ruminococcus gnavus* | Leibniz Institute DSMZ - German Collection of Microorganisms and Cell Cultures GmbH |
| MCC3901 | *Ruminococcus gnavus* | Etrolizumab clinical trial; UC patient; stool<sup>3,4</sup> |
| MCC3902 | *Ruminococcus gnavus* | Etrolizumab clinical trial; UC patient; stool<sup>3,4</sup> |
| MCC3903 | *Ruminococcus gnavus* | Etrolizumab clinical trial; CD patient; stool<sup>3,4</sup> |
| MCC5206 | *Ruminococcus gnavus* | Etrolizumab clinical trial; CD patient; stool<sup>3,4</sup> |
| MCC5209 | *Ruminococcus gnavus* | Etrolizumab clinical trial; CD patient; stool<sup>3,4</sup> |
| MCC5216 | *Ruminococcus gnavus* | Etrolizumab clinical trial; UC patient; stool<sup>3,4</sup> |
| MCC5217 | *Ruminococcus gnavus* | Etrolizumab clinical trial; CD patient; stool<sup>3,4</sup> |
| RJX1126 | *Ruminococcus gnavus* | Laboratory of Ramnik Xavier<sup>5</sup> |
| RJX1128 | *Ruminococcus gnavus* | Laboratory of Ramnik Xavier<sup>5</sup> |

## Supplementary Figures

**Supplementary Figure 1.**
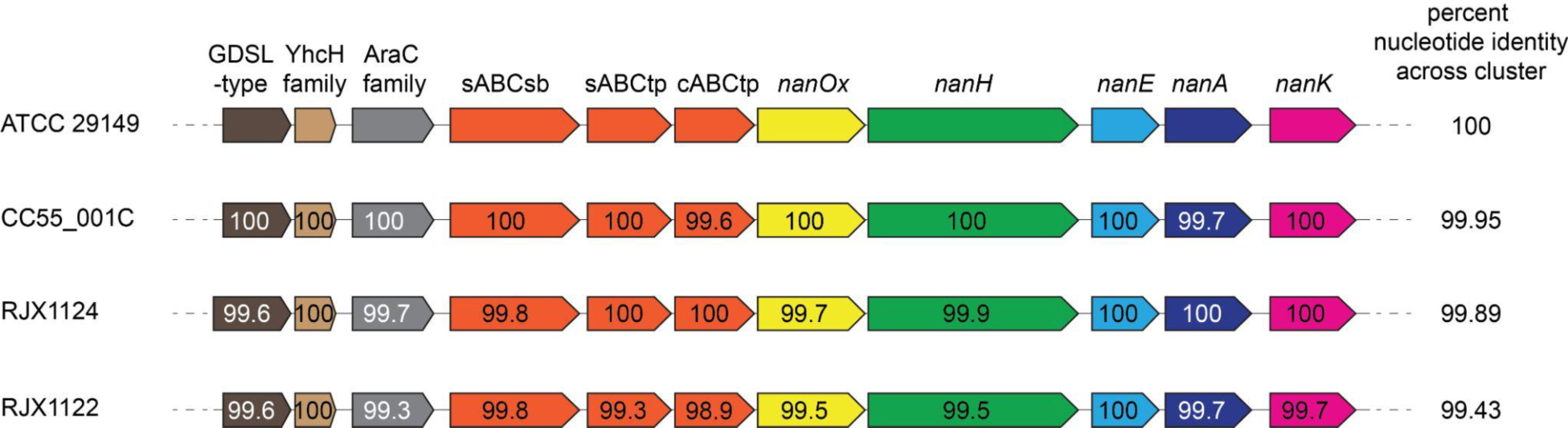
Synteny and sequence identity of the *nan* cluster across diverse *R. gnavus* strains. Four strains sampling the phylogenetic diversity of the *R. gnavus* clade were selected for this analysis. The overall nucleotide identity of this cluster relative to ATCC 29149 is recorded on the right. The numbers inside of each open reading frame indicate the percent identity of each encoded protein at the amino acid level, relative to ATCC 29149.

**Supplementary Figure 2.**
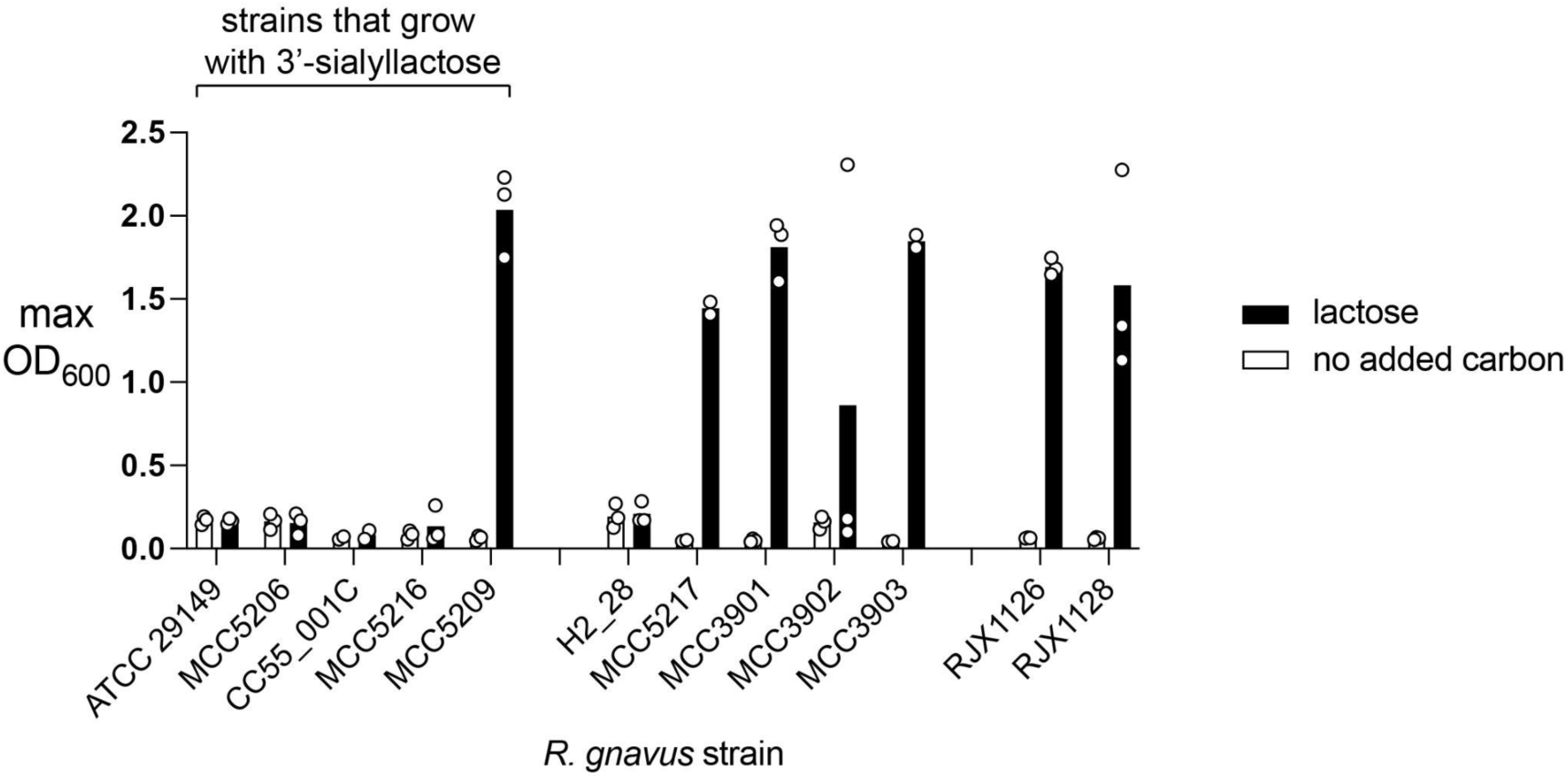
*R. gnavus* strain growth in basal YCFA medium supplemented with lactose. Bars indicate the average of at least two independent experiments, each represented by a single data point.

**Supplementary Figure 3.**
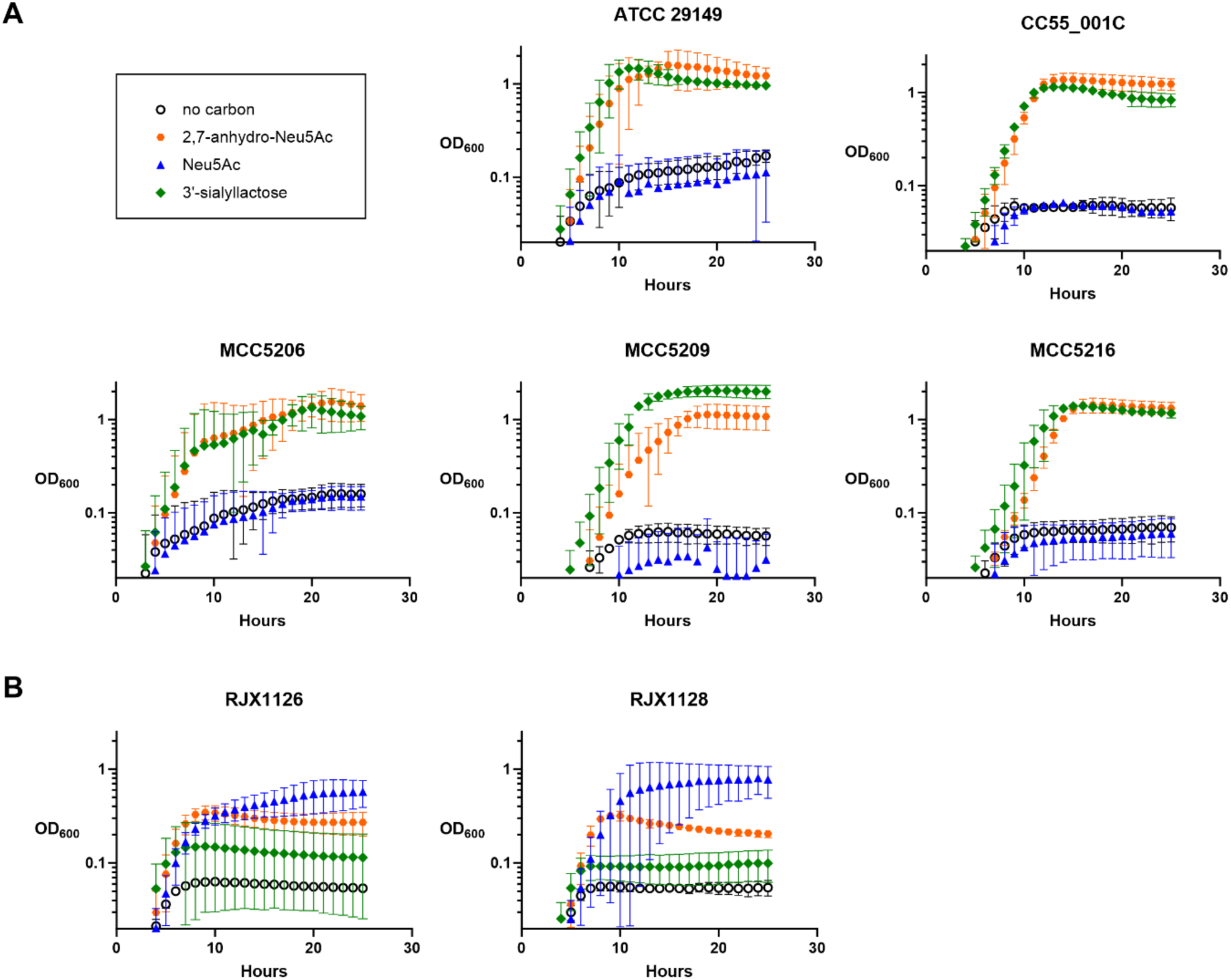
*R. gnavus* strain growth in basal YCFA medium supplemented with simple sugars. (A) 2,7-anhydro-Neu5Ac metabolizing strains, (B) Neu5Ac catabolizing strains. Data points and error bars indicate the average and standard deviation, respectively, of at least two independent experiments

**Supplementary Figure 4.**
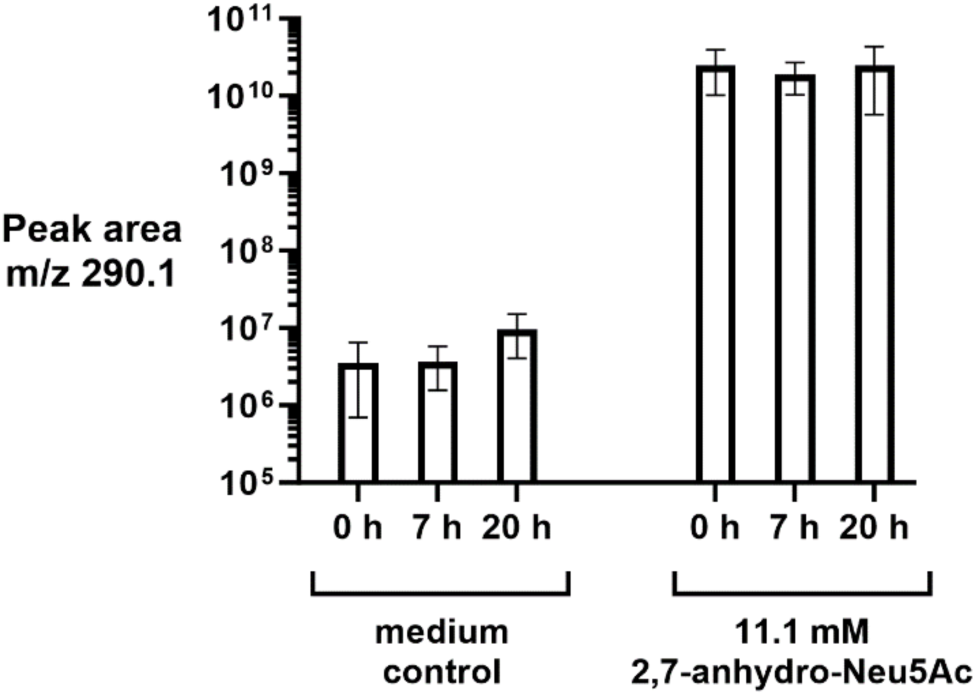
Stability of 2,7-anhydro-Neu5Ac in bYCFA medium at 37 degrees, as demonstrated by liquid chromatography-mass spectrometry (LC-MS)-based detection of 2,7-anhydro-Neu5Ac in the medium at various time points. Bars and error brackets represent the mean and standard deviation, respectively, of three technical replicates. For method details, see Supplementary Materials and Methods.

**Supplementary Figure 5.**
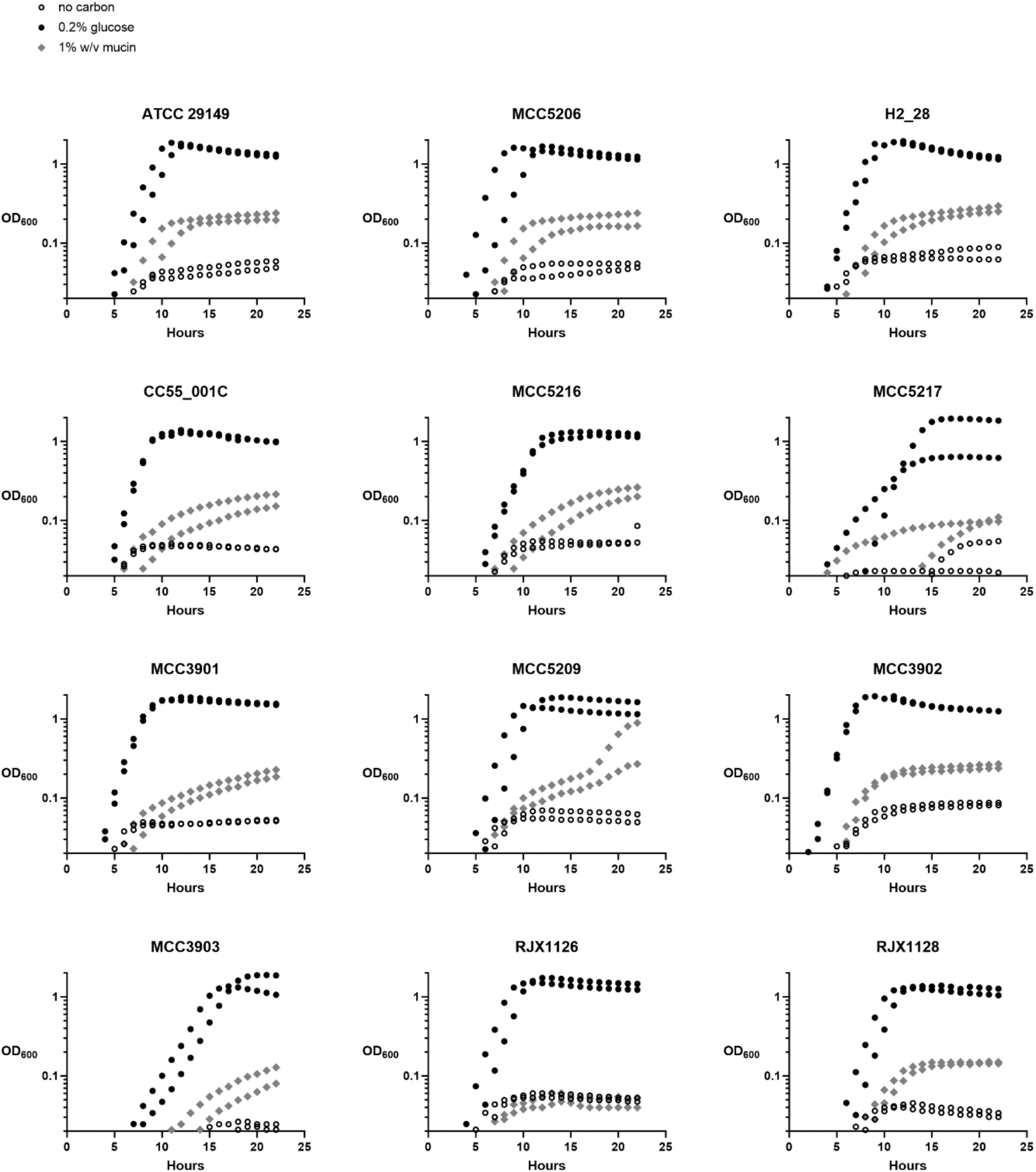
*R. gnavus* strain growth in basal YCFA medium supplemented with purified porcine gastric mucin (pPGM). Data points represent two independent experiments.

## Supplementary Materials and Methods

### gBlock sequence for assembling NanH expression vector

The following gBlock encoding *R. gnavus* NanH, codon-optimized for *E. coli*, was cloned into pET11a pre-digested with NdeI/BamHI, by Gibson assembly:

GTTTAACTTTAAGAAGGAGATATACATATGGCACACCATCATCACCATCATAGTAGCGGCCTGGAAAACTTATACT

**NdeI 6x His TEV cleavage site**

TCCAAGGCTCGAGCCAAGAAGCGCAAACAGACGTGATAGAAGCAGTTGCAGAGAAAAAGCAAGATACAGAGAG

5’ *nanH* coding sequence (no signal sequence, residues 1-25)

TTCCTCTGTTCCTGTCTTGCAGAAAGAAGGGATTGAAATCTCAGAGGGTACCGGATACGATTTGTCAAAAGAGCCGGGGGCCGCCACTGTTAAGGCATTAGAACAGGGCACCATCGTGATCTCATATAAGACAACGTCCGAGAATGCCATACAATCACTTCTTTCGGTGGGTAATGGTACGAAGGGGAACCAGGATAGACACTTTCATCTTTACATAACAAATGCTGGCGGAGTCGGTATGGAATTGAGAAACACCGATGGCGAGTTTAAGTATACCCTGGACTGCCCTGCTGCGGTTCGTGGATCTTATAAGGGGGAACGTGTCTCCAATACGGTGGCACTTAAGGCGGACAAGGAGAATAAACAGTACAAATTATTCGCCAACGGTGAACTTATAGCGACGTTGGACCAAGAAGCATTCAAATTCATAAGCGATATAACCGGAGTAGATAACGTAATGCTGGGAGGCACTATGCGGCAAGGAACAGTGGCCTACCCCTTCGGTGGGTCCATAGAGCGGATGCAAGTCTATCGGGATGTGTTGTCGGACGACGAGCTTATTGCCGTTACAGGCAAAACAATTTATGCTGAAAATATCTTTTACGCTGGGGATGCCACGAAAAGTAATTATTTTAGAATTCCGAGCTTGTTAGCGCTTGACTCCGGCACCGTGATCGCGGCCGCGGACGCACGCTACGGTGGGACACACGATGCGAAGTCGAAAATCAATACGGCCTTTGCAAAAAGCACAGATGGAGGGAAAACATGGGGGCAGCCCACTTTACCATTGAAATTCGATGACTATGTAGCAAAGAACATTGACTGGCCGCGCGATAGCGTAGGAAAAAACGTGCAAATCCAGGGAAGTGCATCTTACATAGACCCTGTACTTCTGGAAGATAAGGAGACACACCGGGTATTCCTGTTCGCAGATATGATGCCCGCAGGCATTGGAAGTTCAAATGCTTCGGTCGGGTCTGGGTTTAAAGAGGTCGATGGTAAGAAATACCTTAAGCTTCACTGGAAGGACGATGCTGCGGGTACGTACGATTACAGCGTGCGGGAAAACGGGACTATTTATAACGATACGACAAACTCCGCGACTGAGTATTCCGTGGACGGGGAATATAATCTTTACAAAAATGGTAATGCAATGTTATGCAAACAATATGATTACAACTTTGAGGGAACCAAATTGCTGGAGACTCAAACTGATACAGATGTCAATATGAACGTGTTCTACAAAGACGCAGACTTTAAAGTATTTCCAACTACCTACTTGGCCATGAAGTATTCCGACGACGAAGGAGAAACGTGGAGTGATCTGCAAATAGTCTCGACCTTCAAGCCGGAGGAAAGCAAGTTTCTTGTATTAGGACCTGGAGTAGGGAAACAAATAGCGAATGGAGAGCATGCCGGTAGACTGATAGTGCCCTTGTATTCGAAATCAAGTGCCGAGTTGGGTTTCATGTACTCGGATGATCACGGTAATAATTGGACGTATGTAGAAGCAGACCAGAATACCGGCGGCGCCACCGCCGAAGCGCAAATCGTGGAAATGCCCGATGGATCGTTGAAAACCTACTTGCGTACAGGGAGTGGGTACATAGCTCAGGTAATGAGTACTGACGGTGGTGAGACATGGTCCGAACGGGTTCCTCTGACGGAGATCGCCACAACAGGTTATGGCACACAGCTTTCAGTCATCAACTACTCTCAGCCGGTGGATGGAAAACCAGCAATATTACTGAGTGCGCCTAATGCGACGAACGGACGGAAGAATGGGAAAATTTGGATTGGATTAATAAGCGAAACTGGCAACTCTGGAAAGGATAAATACTCTGTGGACTGGAAGTACTGCTATTCCGTCGACACGCCGCAGATGGGTTATAGCTATTCTTGTCTTACCGAATTGCCGGATGGAGAAATAGGCCTTCTTTATGAGAAATATGATAGTTGGTCACGGAACGAATTACATTTAAAAAACATTCTTAAGTATGAGCGCTTTAATATTGATGAGCTTAAAGTACAACCTTAAGGATCCGGCTGCTAACAA

**nanH - 3’ end BamHI**

### Chemical analysis of 2,7-anhydro-Neu5Ac synthesis product

Liquid chromatography-mass spectrometry (LC-MS) analysis was performed using an Agilent 1290 ultra-high-performance liquid chromatography system coupled to an Agilent MSD (6140) mass spectrometer, using electrospray ionization. LC separation was performed using a Phenomenex XB-C18 column (1.7μm, 50 × 2.1 mm), at 40 °C and a flow rate of 0.4 mL/min. Mobile phase A (MPA) was water with 0.1% FA and mobile phase B (MPB) was acetonitrile with 0.1% FA. The LC method consisted of an initial equilibration at 2% MPB for 1.5 min, followed by a gradient from 2 - 98% MPB over 7 min, and terminating in a hold at 98% MPB for 1.5 min. UV absorbance was collected at 220 nm and 254 nm and a full mass spectrometry scan was applied to all experiments. 2,7-anhydro-Neu5Ac m/z = 292.1 [M+H]^+^.

NMR spectra were recorded on a Bruker Avance 500 MHz spectrometer with an HCN cryoprobe. Data were acquired at 298 K in DMSO-d6, with chemical shifts referenced to 2.50 ppm in ^1^H and 39.51 ppm in ^13^C; or at 298 K in D_2_O, with chemical shifts referenced to 4.79 ppm in ^1^H. Chemical shifts are expressed in δ ppm. ^1^H NMR data: (500 MHz, D_2_O) δ 4.55 (s, ^1^H), 4.45 (d, J = 7.8 Hz, ^1^H), 3.96 (d, J = 5.5 Hz, ^1^H), 3.93 (s, ^1^H), 3.77 (dd, J = 11.9, 3.1 Hz, ^1^H), 3.61 (dd, J = 12.0, 6.1 Hz, ^1^H), 3.55 (td, J = 6.8, 2.9 Hz, ^1^H), 2.18 (dd, J = 15.3, 5.6 Hz, ^1^H), 2.05 (s, ^3^H), 2.02 (d, J = 15.7 Hz, ^1^H).

### LC-MS analysis of 2,7-anhydro-Neu5Ac stability in culture medium

Media samples were quenched with 8:2 methanol:water, sonicated, and centrifuged for protein crush and precipitation. Supernatants were dried down by N_2_ gas and reconstituted in 8:2 acetonitrile:water. LC-MS analysis was carried out using a Shimazu LC system coupled to a Thermo QEplus Obitrap Mass Spectrometer. For LC separation, a Waters BEH Amide column (2.1 mm × 100 mm, 1.7 µm) was used with mobile phase A of water and mobile phase B of 95:5 ACN:water, both buffered with 10 mM ammonium formate and 0.125% formic acid. LC was performed at a flow rate of 0.4 mL/min and 40 °C with following gradient: 0 min 100% B, 0.5 min 100% B, 2 min 70 B%, 7.7 min 40% B, 9.5 min 30% B, 10.25 min 30% B, 12.75 min 100 % B, 16.75 min 100% B. MS was operated in negative ion ddMS2 mode to acquire accurate mass, retention time, and MS/MS at a mass accuracy of 5 ppm and mass resolution of 70000.

